# Aging leads to sex-dependent effects on pair bonding and increased number of oxytocin-producing neurons in monogamous prairie voles

**DOI:** 10.1101/2024.05.17.594752

**Authors:** Michael A. Kelberman, Kelly E. Winther, Yana M. Medvedeva, Zoe R. Donaldson

## Abstract

Pair bonds powerfully modulate health, which becomes particularly important when facing the detrimental effects of aging. To examine the impact of aging on relationship formation and response to loss, we examined behavior in 6-, 12-, and 18-month male and female prairie voles, a monogamous species that forms mating-based pair bonds. We found that older males (18-months) bonded quicker than younger voles, while similarly aged female voles increased partner directed affiliative behaviors. Supporting sex differences in bonding behaviors, we found that males were more likely to sample both partner and novel voles while females were more likely to display partner preference during the initial 20 minutes of the test. Using partner separation to study loss, we observed an erosion of partner preference only in 12-month females, but an overall decrease in partner-directed affiliation in females across all groups, but not in males. Finally, we found that the number of oxytocin, but not vasopressin, cells in the paraventricular hypothalamus increased during aging. These results establish prairie voles as a novel model to study the effects of normal and abnormal aging on pair bonding.

**Highlights:**

- 18-month male voles demonstrate accelerated bond formation
- 18-month female voles increase partner-directed huddling after 2 wks
- Bonds erode faster in 12-month female voles after partner separation
- Female behavior from partner preference tests is reflected in free interaction
- The number of paraventricular hypothalamus oxytocin cells increase during aging

## 1. Introduction

Human social behaviors are complex, evolve over a life course, and become dysfunctional in age-related disorders, such as Alzheimer’s disease (AD). During normal aging, social relationships become more positive and satisfying with age, and social engagement contributes to cognitive resilience, even in the presence of AD pathology (Charles and Piazza, 2007; Luong et al., 2011; Scarmeas et al., 2001; Stern, 2012). Meanwhile, the onset of irritability, depression, heightened anxiety fracture social relationships in prodromal phases of AD and deteriorating social memory in mid to late stages of disease exacerbate relationship stress, negatively impacting the emotional well-being of AD patients and their friends and family (Bediou et al., 2009; Desmarais et al., 2018; Ehrenberg et al., 2023, 2018; Müller-Spahn, 2003; Ren et al., 2023). However, our preclinical understanding of social changes over the course of normal aging, especially into geriatric periods, remains limited to mice and rats, which do not display many of the complex social phenotypes fundamental to humans.

Monogamous prairie voles (*Microtus ochrogaster*) provide a promising model for studying the effects of normal and abnormal aging on complex sociocognitive processes relevant to humans. Prairie voles, unlike mice and rats, form life-long pair bonds and have a lifecourse that is similar although not identical to that of laboratory mice and rats. They have a ∼21 day gestation and can wean at 20 days. However, they display somewhat faster development with earlier tooth eruption, eye opening, fur growth, accelerated patterns of cortical gene expression for some genes (James et al., 2022; Shapiro and Insel, 1990; Spangenberg et al., 2014), and are productively active at an earlier age (females can get pregnant as early as post-natal day 40 if housed with a male; Solomon, 1991). They display remarkable variation in longevity in the wild and in lab where they have been documented to reach anywhere from 18 months to 5 years (Fischer, 1945; Getz et al., 1997; Grippo et al., 2021; Kenkel et al., 2019; Stalling, 1990).

Pair bonding – like other species-typical social behaviors – is mediated by the neurohormones, oxytocin and vasopressin. These peptide hormones are produced primarly in the paraventricular hypothalamus (PVH) and supraoptical nucleus, with the former contributing to most neural release. Differences in nonapeptide receptor densities in key brain areas such as the nucleus accumbens, prelimbic cortex, and ventral pallidum account, in part, for species differences in pair bonding (Insel and Shapiro, 1992; Olazábal and Young, 2006; Ophir et al., 2012; Young et al., 2001, 1996). Furthermore, direct manipulation of these circuits can facilitate or inhibit pair bonding in prairie voles, and can even produce bonding in typically promiscuous species (Keebaugh et al., 2015; Lim et al., 2004; Lim and Young, 2004; Ross et al., 2009; Winslow et al., 1993; Young et al., 2001). Dynamic changes in oxytocin and vasopressin circuitry occur across postnatal development and in response to pair bonding, pup rearing, and partner separation (Audunsdottir and Quintana, 2022; Ebner et al., 2013; Fliers et al., 1985; Fricker et al., 2023; Hiura et al., 2023; Hiura and Ophir, 2018; Ishunina and Swaab, 1999; Kelly et al., 2018, 2017; Kenkel et al., 2019). However, the vast majority of experimentation on pair bonding and its component neurochemical systems is performed in young adult voles.

The dynamics of pair bonding have been well studied in younger animals where female animals form a partner preference more rapidly than male animals (Brusman et al., 2021; DeVries and Carter, 1999; Harbert et al., 2020; Insel and Hulihan, 1995). However, the impacts of age, and its potential interaction with sex, have been studied only in restricted settings. For instance, 2-3 years old male voles show a reduction in the amount of time spent huddling with a stranger compared to younger males (Kenkel et al., 2019), and ∼4-year-old male voles display increased immobility during forced swim following separation from their long term partners (Grippo et al., 2021). Yet neither of these studies examined female behavior. Other studies have looked at social behavior outside the context of pair bonds, observing reductions in both prosocial behaviors directed towards a familiar same-sex sibling and aggression towards stranger conspecifics of either sex in non-bonded voles up to ∼9 months (∼mid-adulthood; Powell et al., 2022). Thus, work to date supports the hypothesis that there are sex- and age-dependent changes in social behavior, bonding, and their underlying hormonal bases, but it has not been thoroughly evaluated.

To comprehensively delineate the age- and sex-dependent changes in social behavior and oxytocin/vasopressin, we tested 6-, 12-, and 18-month-old naive prairie voles in partner preference tests (PPTs) and during unconstrained free interaction. These ages were chosen as representative of early (6 mo) and late (12 mo) adulthood, and into the “geriatric” (18 mo) period for voles. Animals were tested 2 days (short-term) and 2 wks (long-term) following pairing to examine whether age impacts the development of pair bonds (Brusman et al., 2021). We then separated pairs for 4 weeks before testing for bond persistence following partner loss via partner preference and free interaction tests (Fricker et al., 2023; Sadino et al., 2021). Finally, we investigated the potential neurochemical mechanisms underlying the evolution of pair bonds dynamics during aging by quantifying the number of oxytocin and vasopressin cells in the PVH. Our results indicate sex-dependent effects of age on pair bonding such that old males bond quicker while females increase partner-directed affiliative behavior and undergo a U-shaped curve of partner preference following long-term partner loss. These trends were evident within the first 20 minutes of the PPTs, and female behavior during PPTs was further reflected during unconstrained free interaction, suggesting that females behavior dictates pairwise interactions (Brusman et al., 2021). These changes were met by age-dependent increases in the number of oxytocin, but not vasopressin, neurons within the PVH. Overall, these data provide fundamental knowledge on pair bonding during normal aging in prairie voles which can be used to further develop the species for studying the bidirectional interplay between abnormal aging and socioemotional wellbeing.

## 2. Materials and methods

### 2.1 Animals

Prairie voles were bred in-house at the University of Colorado Boulder in a colony originating from wild caught animals from Illinois. Animals were housed in same-sex groups of 2-4 animals in static rodent cages until pairing (see section *2.2. Experimental Timeline*), with free access to water, rabbit chow supplemented with sunflower seeds, and cotton nestlets, igloos, and tubes for enrichment. Housing conditions were maintained at 23-26°C on a 14:10 hr light:dark cycle. All procedures were performed during the light cycle and approved by the Institutional Animal Care and Use committee at the University of Colorado Boulder.

### 2.2 Experimental Timeline

Experiments were performed on opposite-sex, non-sibling pairs at 6 (N = 10 pairs; mean age at pairing: 6.2 months, age range at pairing: 5.5-6.4 months), 12 (N = 10 pairs; mean age at pairing: 11.9 months, age range at pairing: 11.3-12.4 months), or 18 months (N = 7 pairs; mean age at pairing: 18.6 months, age range at pairing: 17.7-21.4 months). Animals were paired (Day 0) and partner preference tests were performed on both members of the pair following 2 days (short-term) and 16 days (long-term/2 wks) of pairing and cohabitation, pairs were subjected to partner preference tests. On days 3 and 17, animals were tested on a free interaction test. After completing the free interaction test on day 17, pairs were separated and single housed for 4 weeks in separate vivarium rooms. Animals were subjected to a final partner preference test on day 44 and free interaction on day 45.

### 2.3 Partner Preference Tests

Testing was performed as described previously (Brusman et al., 2021; Sadino et al., 2021). Partner and stranger, non-sibling animals were tethered to the ends of a three-chamber plexiglass arena (76.0 cm long, 20.0 cm wide, and 30.0 cm tall) following brief anesthetization with isoflurane. Tethers were made of an eyebolt attached to a chain of fishing swivels, with animals being connected via zip ties around the animal’s neck. All animals had access to food and water for the duration of the test (3 hours). Experimental animals were placed into the center chamber separated from the other two chambers by opaque dividers. The opaque dividers were removed, and the experimental animal was allowed to move freely around the arena for 3 hrs. Tracking was performed by overhead Panasonic WVCP304 cameras to record eight chambers simultaneously. Movement from all animals were scored using TopScan software according to (Ahern et al., 2009; Brusman et al., 2021). Both voles in a pair were tested back-to-back on each test days, separated by approximately 1 hr. Order of testing and side of partner was randomized and counterbalanced within an age group. The main outcome metric of these tests were time spent huddling with partner and partner preference (partner huddle time/[partner huddle time + stranger huddle time]). We also asked whether any partner preference phenotypes were a result of which animal (partner versus stranger) or area of the PPT chambers (partner side, stranger side, or center) that the focal animals chose to spend their time, which we included as additional metrics from PPTs.

We decided to investigate response and habituation to social novelty during PPTs, and their dependence on age and test day. We therefore analyzed the first 5 minutes and the following 15 minutes of each PPT. During this analysis one male animal from the 12-month cohort did not leave the center chamber of the arena within the first 20 minutes of the 2 wk PPT. Similarly two female animals from the 6-month age group were excluded in these analyses for failing to leave the center chamber during the initial 20 minutes of the 4 wk separation PPT. For this analysis we calculated an acute preference index ([partner huddle time – stranger huddle time]/[partner huddle time + stranger huddle time]).

### 2.4 Free Interaction

24 hrs after each partner preference test, pairs were allowed to freely interact in a plexiglass arena (50.7 cm long, 20.0 cm wide, and 30.0 cm tall), as previously described (Brusman et al., 2021). Animals in a pair were placed on opposite sides of the two-chamber arena separated by an opaque divider. The divider was removed, and animals were allowed to freely explore the chamber for 3 hrs. Overhead Logitech C925e webcams were used to record four free interaction tests simultaneously. Social contact between the two animals were scored using TopScan High-Throughput software v3.0 (Cleversys Inc) using scoring methods optimized within our lab (Ahern et al., 2009; Brusman et al., 2021). Identity swapping was common during unconstrained free interaction, so the main metric we retained for analysis was time spent in close social contact (defined by setting the “joint motion” parameter to <5).

### 2.5 PVH Oxytocin and Vasopressin Staining, Imaging, and Cell Counts

24-72 hrs following the final free interaction test, a subset of animals from each age were injected with a mixture of ketamine/xylazine and perfused with 1x phosphate buffered saline (PBS) followed by 4% paraformaldehyde. Brains were stored in 4% paraformaldehyde for 24-48 hrs and then transferred to 30% sucrose. Brains were sliced at 40µm on a Leica SM2010R microtome and slices were stored in 1x PBS with 0.01% sodium azide at 4°C until staining. 1-3 sections for each animal encompassing the PVH were atlas matched (approximately -0.94 to -1.7 from bregma) to the Mouse Brain in Stereotaxic Coordinates Compact Third Edition. Sections were washed 3 x 5 minutes in 1x PBS and then incubated for 1 hr at room temperature in blocking solution (0.2% Triton-X, 6% bovine serum albumin, and 10% normal goat serum in 1x PBS). Sections were then incubated in 1:500 primary antibodies in blocking solution, mouse anti-NeuN (abcam, ab104224), rabbit anti-oxytocin (Immunostar, 20068), and guinea pig anti-vasopressin (BMA Biomedicals, T-5048), for 48 hrs. After 48 hrs, sections were washed 3 x 5 min in 1x PBS and then incubated in 1:500 Alexa Fluor™ secondary antibodies, goat anti-mouse 405 (ThermoFisher Scientific, A-31553), rabbit 488 (ThermoFisher Scientific, A-11008), and guinea pig 568 (ThermoFisher Scientific, A-11075), diluted in blocking solution for 2 hrs. Sections were wash 3 x 5 min in 1x PBS and mounted on Diamond White Glass microscope slides (Globe Scientific, INC.), coverslipped with ProLong™ Diamond Antifade Mountant (ThermoFisher Scientific, P36970), and sealed with nail polish.

Images were acquired on an Olympus iX83 wide field slide scanner with a Chroma Multi LED filter set (Cat: 69401) at 20x for quantification with the following consistent exposures times (DAPI 300ms, FITC 50ms, TRITC 100ms). Representative images for figures were acquired at 10x. Images used for quantification were imported into ImageJ/Fiji and rolling ball subtraction was used to reduce image background (oxytocin = 100, vasopressin = 750). Images were grayscaled and thresholded using the Otsu method (oxytocin = 0-45, vasopressin = 0-65). The watershed function was applied to separate individual cells and analyze particles (size = 50-infinity, circularity = 0.3-1.00) was performed to count the number of oxytocin and vasopressin cells in a region of interest with a standard size. Counts were averaged across all sections for an animal. Males and female histology results were combined due to the smaller sample size; both sexes showed the same overall trends.

### 2.6 Statistical Analysis and Data Visualization

Data analysis and visualization was performed in GraphPad Prism (version 10.0.0). All behavioral data were analyzed as a two-way repeated measures ANOVA considering age and test day as factors. When there were interactions between age and test day, we followed up with Tukey’s post-hoc tests. To determine whether groups demonstrated a significant partner preference, we utilized a one-sample t-test to compare group means to the null hypothesis of 0.5 (no preference for partner or stranger). A similar method was used for acute preference indices but compared to the null hypothesis of 0 (no preference for partner or stranger). Histology data was analyzed with a one-way ANOVA considering age as a factor and significant main effects of age were followed up with Tukey’s post-hoc tests.

## 3. Results

### 3.1 Aging accelerates bond formation in male prairie voles

We first asked whether there were any differences in behavioral metrics during PPTs that depend on age. Because there are sex differences in the development of bonds (Brusman et al., 2021; DeVries and Carter, 1999; Insel and Hulihan, 1995), we analyzed males and females separately. An experimental timeline for PPTs can be found in **Figure 1A**. We first examined whether there were any differences in partner preference score following 2 days and 2 wks cohabitation, or after 4 wks separation in male animals (**Figure 1B**). There were no effects of age, test day or an age x test day interaction on partner preference score (all p>0.05). We also performed a one-sample t-test against the null hypothesis of no preference (50%), which revealed that 18-month males formed a strong partner preference after only 2 days of cohabitation (t_6_=20.32, p<0.0001). In contrast, 6- and 12-month male animals developed a significant partner preference only after 2 wks of cohabitation (6 month: t_9_=3.596, p=0.0058; 12 month: t_9_=4.929, p=0.0008), which was also maintained by 18-month males (t_6_=14.25, p<0.0001). Males across all ages maintained their partner preference after 4 wks separation from their partners (6 month: t_9_=4.555, p=0.0014; 12 month: t_9_=3.382, p=0.0081; 18 month: t_6_=17.2, p<0.0001). There were no differences in the amount of time male animals spent huddling with partner or stranger animals as a result of age, test day, or an age x test day interaction (all p>0.05; **Figure 1C and D**). Similarly, there was no effect of age, time, or an age x time interaction on proportion of time spent in the partner, stranger, or center chamber of the PPT arena (all p>0.05; **Supplemental Figure 1A-C**).

**Figure 1.**
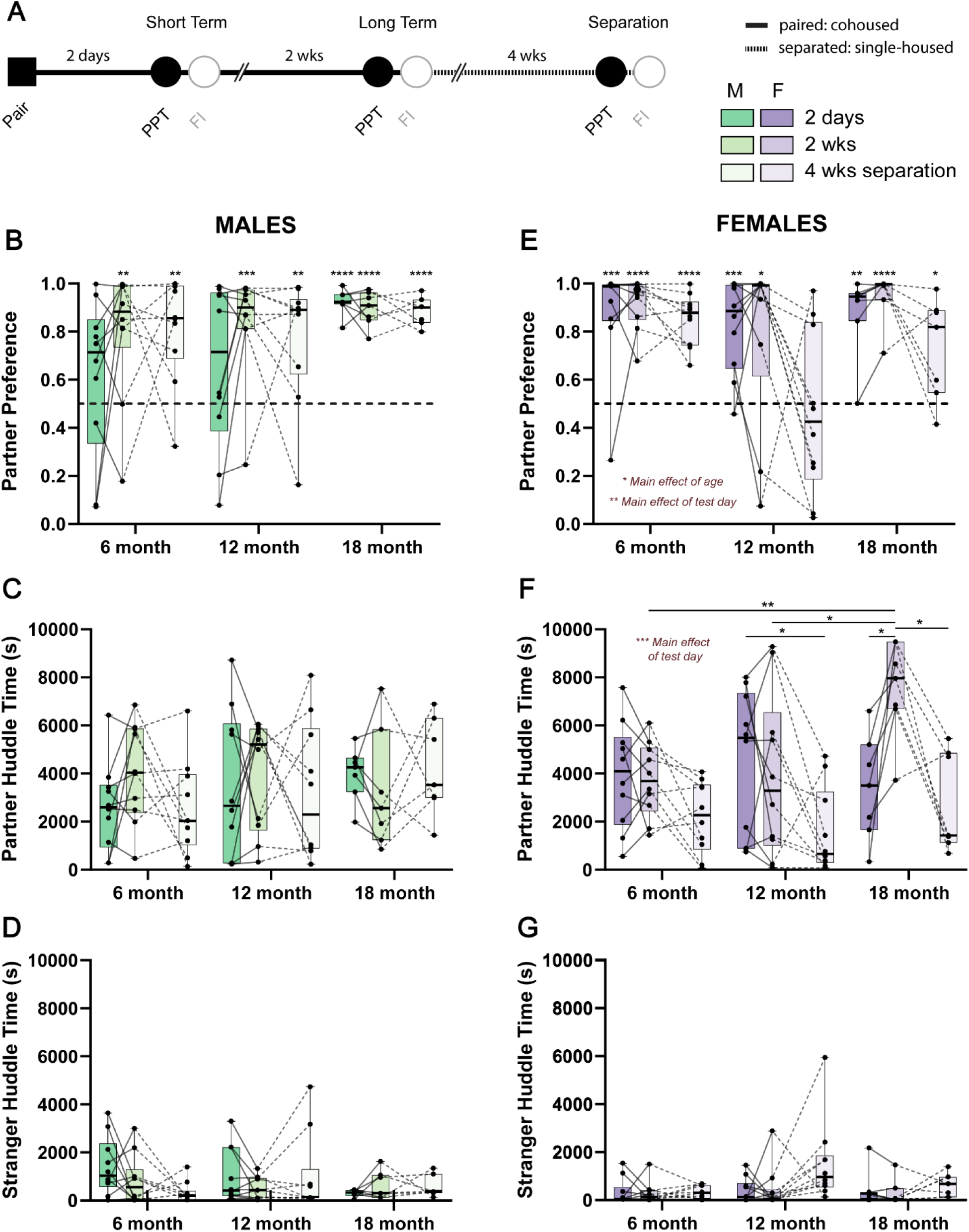
Sex-dependent effects of age on pair bonding. A) Schematic of behavioral timeline where opposite sex voles were paired before undergoing partner preference tests on day 2 (short-term) and 16 (long-term/2 wks). Pairs were then separated for 4 wks and then tested on another partner preference test. PPT: partner preference test, FI: free interaction. B) Partner preference scores in male 6-, 12-, and 18-month voles. 6- and 12-month male voles only form a partner preference after 2 wks of cohabitation and maintain this preference after 4 wks separation. Meanwhile 18-month male voles form a robust partner preference after 2 days cohabitation which is maintained on other test days. Partner preference was tested against the null hypothesis of no preference (0.5, dotted line). C) Partner huddle time is no different in male animals due to age or test day. D) Similarly, there are no differences in stranger huddle time due to age or test day in male animals. E) Partner preference scores in 6-, 12-, and 18-month female animals. Female animals regardless of age and test day displayed a significant partner preference (compared to the null hypothesis of no preference of 0.5) F) Unlike males, 18-month females increased their time spent huddling with their partner, specifically after 2 wks of cohabitation, which wasn’t apparent in other age groups. Generally, females in all age groups decreased partner huddle time after 4 wks separation. G) Like males, there were no effects of age or test day on time spend huddling with stranger animals. N=7-10 animals per group; *p<0.05, **p<0.01, ***p<0.001 ****p<0.0001.

### 3.2 Age affects affiliative behavior and partner preference after loss in female prairie voles

We next examined whether there were differences in behavioral phenotypes during PPTs in female animals (**Figure 1E-G**). Similar to previous reports, female animals formed a partner preference following 2 days cohabitation regardless of age (6 month: t_9_=5.341, p=0.0005; 12 month: t_9_=5.293, p=0.0005; 18 month: t_6_=5.794, p=0.0012), which was maintained after 2 wks (6 month: t_9_=12.56, p<0.0001; 12 month: t_9_=2.645, p=0.0267; 18 month: t_6_=11.05, p<0.0001). Partner preference was also conserved following 4 wks separation, but only in 6- and 18-month animals (6 month: t_9_=10.4, p<0.0001; 18 month: t_6_=2.904, p=0.0272). Moreover, there was a significant effect of age (F_2,24_=4.686, p=0.0191) and time (F_1.688,40.5_=7.663, p=0.0025), but not an age x time interaction (F_4,48_=1.894, p=0.1268) on partner preference scores. In general, 12-month females displayed lower partner preference scores across each test day compared to other age groups, which was most apparent after 4 wks separation. In addition, females showed lower partner preference scores after 4 wks separation compared to the 2 days and 2 wks timepoints.

Unlike males, there were also effects of test day (F_1.956,46.95_=13.24, p<0.0001) and an age x test day interaction (F_4,48_=3.419, p=0.0154) on the amount of time females spent huddling with their partners (**Figure 1F**). Females huddled more with their partners after 2 days and 2 wks of cohabitation compared to after 4 wks separation. Post-hoc tests revealed that specifically 18-month female animals displayed increased partner directed huddling at 2 wks compared to both 2 days (q_6_=5.71, p=0.0161) and 4 wks separation (q_6_=5.76, p=0.0154). 18-month females also displayed a greater amount of partner directed huddling compared to both 6-(q_10.71_=5.93, p=0.004) and 12-month (q_14.75_=3.91, p=0.04) females after 2 wks. Partner directed huddling time was lower following 4 wks separation compared to 2 days only in the 12-month females (q_9_=3.969, p=0.0489).

These phenotypes occurred in the absence of significant differences in stranger huddle time across age and test days (all p>0.05; **Figure 1G**). However, partner directed behaviors were associated with differences in which parts of the PPT chambers that female animals spent their time. (**Supplemental Figure 1D-F**). There was a main effect of test day on the proportion of time that females spent in the partner (F_1.795,43.08_=10.25, p=0.0004) and stranger (F_2,24_=4.528, p=0.0215) chambers. At each age, females spent more time in their partner’s chamber after 2 wks cohabitation compared to either 2 days or after 4 wks separation. In addition, females spent less time in their partner’s chamber after 4 wks separation. These effects were mirrored in the opposite direction with respect to the chamber that held the stranger animal. Finally, there was an effect of age on the amount of time females spent in the center chamber (F_2,24_=3.844, p=0.0356), where animals spent gradually less time in the center chamber with increasing age.

### 3.3 Sex differences in partner preference are evident early in partner preference tests

While the partner preference test provides an aggregate measure of social choice and partner- and stranger-directed behavior over the course of 3 hours, we also analyzed the first 5 minutes of each test to examine the acute response to a stranger vole across sex and age (**Figure 2**). We also examined behavior in minutes 6 – 20 to examine emergence of preference behavior after the initial 5-minute investigatory period, essentially asking whether partner preference emerges within this early period in an age- or sex-dependent manner. In voles, we observed a consistent stranger preference in the first 5 minutes of the task only for 6-month males after 2 weeks and 6-month females after 2 days of cohabitation (males: t_9_=2.4, p=0.039; females: t_7_=2.669, p=0.032). Of note, this differs from laboratory mice, where a stranger preference is frequently observed within this timeframe (Beery et al., 2018; Crawley et al., 2007; Moy et al., 2004). In males, it took longer for partner preference to emerge; it was never present in the first 5 minutes, and it emerged in minutes 6 – 20 only in 12- and 18-month males (12- and 18-month) that had undergone partner separation (**Figure 2A and B**). In contrast, females often displayed a partner preference even within the first 5 minutes, which remained evident and was strengthened in the succeeding 15 minutes (**Figure 2C and D**). There was an additional significant effect of test day on the partner preference scores during the initial 5 minutes (F_1,22_=5.558, p=0.0277) and subsequent 15 minutes (F_1,22_=5.343, p=0.031) of the PPTs for female animals only. Females displayed higher partner preference scores at the 2 wk time point across all ages. Beyond the dynamics noted above, there were no main effects of age, test day, or their interaction on partner preference within the first 5 minutes or the next 15 minutes of PPTs (all p>0.05).

**Figure 2.**
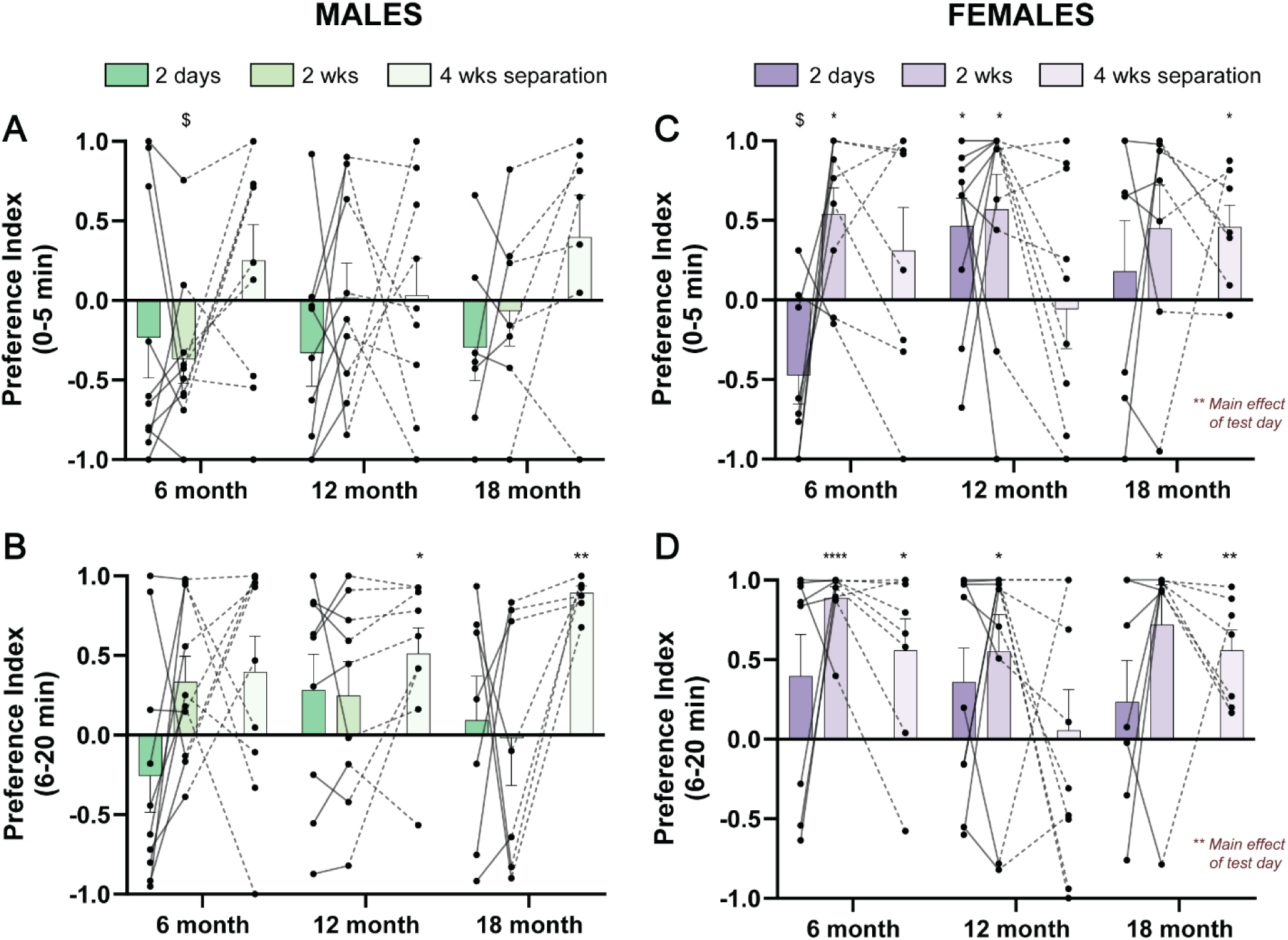
Sex-dependent effects of age on pair bonding are apparent within the initial few minutes of the partner preference test. We used social preference index as a measure of partner or stranger preference during the initial 20 minutes of partner preference tests comparing to a null hypothesis of no preference of 0. A) Male animals displayed a lack of partner or stranger preference during the initial 5 minutes of partner preference tests, with the exception of 6-month animals after 2 wks of cohabitation, which displayed a stranger preference. B) Similarly, most male animals did not display a partner or stranger preference in the following 15 minutes of partner preference tests, except for 12- and 18-month after 4 wks separation, which showed a significant partner preference C) Unlike males, females across ages and test days tended to display a significant partner preference within the initial 5 minutes of partner preference tests, with the exception of 6-month animals after 2 days, which had a stranger preference. D) Female animals of all ages continued to show a significant partner preference in the following 15 minutes of partner preference tests. N = 7-10 pairs per group; *p<0.05, **p<0.01, ****p<0.0001 for denoting significant main effects and partner preference compared to the null hypothesis of 0 (no preference). ^$^p<0.05 for denoting significant stranger preference compared to the null hypothesis of 0 (no preference).

### 3.4 Social contact time during free interaction reflect trends in partner preference tests

PPTs are useful for assessing social choice as to two of the three animals being tethered. However, we have previously shown that behaviors observed during PPTs are only loosely correlated with those observed during unconstrained dyadic interaction (Brusman et al., 2021). We therefore supplemented PPTs with free interaction tests that occurred the following day, recording total interaction time for each pair. The behavioral timeline for free interaction tests can be found in **Figure 3A**. In the 18-month cohort, 2 of the 7 pairs were excluded due to technical issues with behavioral recording. We observed a significant effect of age (F_2,22_=15.62, p=0.0057) and test day (F_1.526,33.58_=11.61, p=0.0048), but no age x test day interaction (F_4,44_=2.20, p=0.08) on the amount of time that pairs spent in close social contact (**Figure 3B**). 18-month animals displayed high amounts of social contact compared to 6- and 12-month animals at each test day, while social contact generally decreased after 4 wks separation which was most apparent in 12-month pairs.

**Figure 3.**
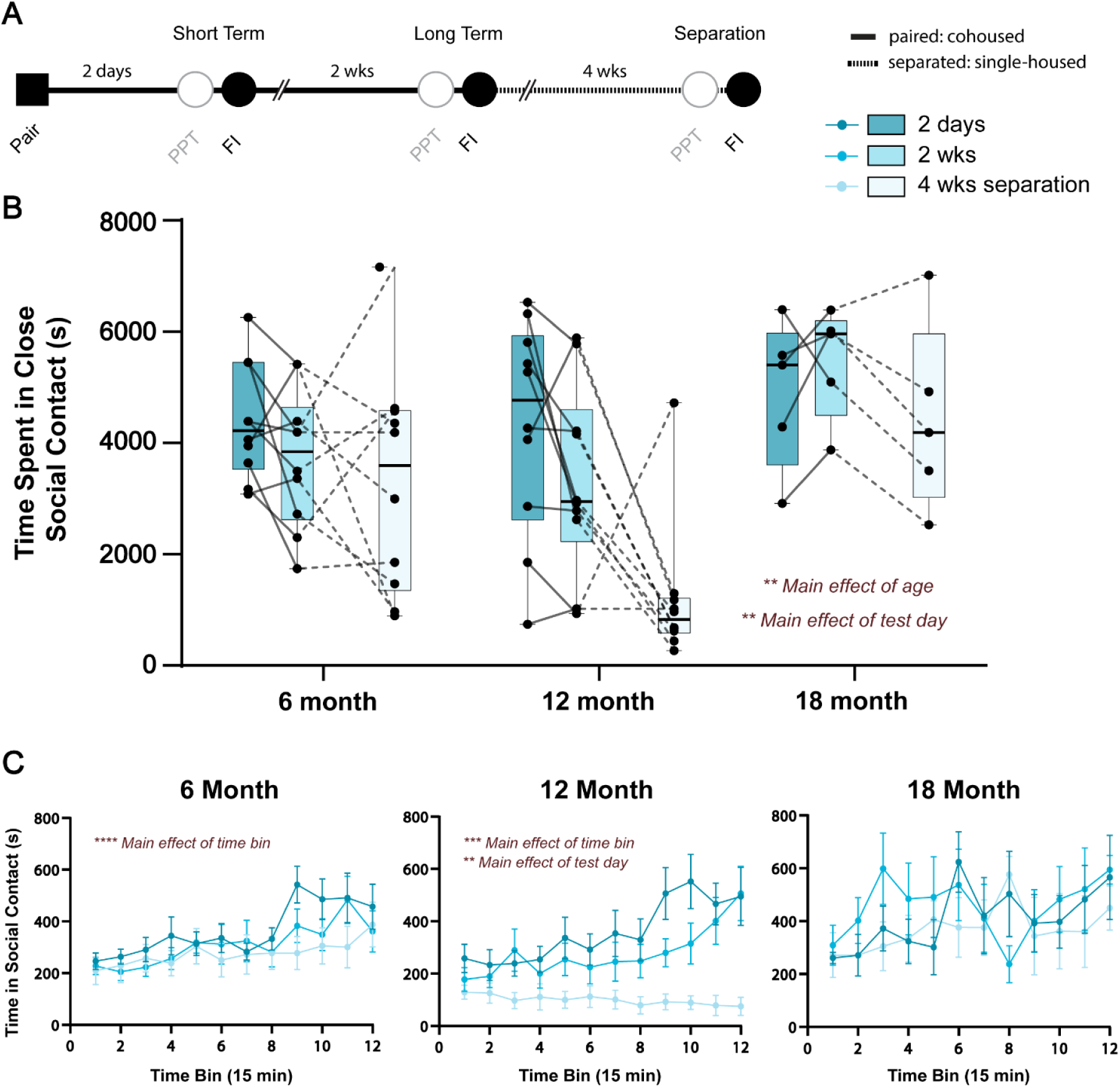
Age and test day impact time spent in close social contact during unconstrained free interaction. A) Schematic of behavioral timeline where opposite sex voles were allowed allowed to interact in an open arena the day after each partner preference test at the short term and long term timepoints and following 4 wks separation. PPT: partner preference test, FI: free interaction. B) 18-month animals tended to spend the most time in close social contact, regardless of test day. Meanwhile, across ages, pairs spent less time in close social contact after 4 wks separation. C) Time spent in close social contact was then separated into 15-minute time bins and analyzed within each age group. There was a main effect of time bin in 6- and 12-month pairs such that time spent in close social contact increased over the course of the 3-hour task. There was an additional main effect of test day in 12-month pairs, where animals spent less time in close social contact after 4 wks separation. An interaction effect between test day and time bin was also apparent for 12-month pairs, and statistics for these comparisons can be found in **Supplemental Table 1**. There were no effects of time bin, test day, or their interaction on the time spent in close social contact during unconstrained free interaction in 18-month pairs. N=5-10 pairs per group; **p<0.01, ***p<0.0001, ****p<0.0001.

To examine social interaction over time, we used 15-minute bins for each age group at each test day (**Figure 3C**). 6- and 12-month animals showed increases in time spent in close social contact, especially within the last hour of the test. A one-way ANOVA demonstrated a main effect of time bin on time spent in social contact in 6-month pairs (F_11,99_=6.757, p<0.0001). Meanwhile 12-month animals displayed a significant effect of time bin (F_11,99_=3.571, p=0.0003), test day (F_2,18_=9.22, p=0.0018), and a time bin x test day interaction (F_22,198_=2.864, p<0.0001) on time spent in close social contact. Notably, time spent huddling generally increased over the course of the 3 hr test, with the exception of the 4 wks separation time point. In addition, huddle time decreased across test days. Post-hoc tests for the 12-month pairs revealed many time bins where huddling time was lower at the 4 wk separation point compared to either other day, in addition to a few timepoints towards the end of the test where huddling time on day 2 was greater than after 2 wks (statistics available in **Supplemental Table 1**). Meanwhile, there were no effects of time bin, test day, or a time bin x test day interaction on time spent in close social contact during free interaction in 18-month animals.

### 3.5 Number of oxytocin, but not vasopressin, cells increase from 12- to 18-months

The nonapeptides oxytocin and vasopressin synthesized in the PVH have been extensively implicated in pair bonding in voles. We used cell counts to gauge whether these systems are altered during the course of aging and could support behavioral changes observed during PPTs and free interaction (**Figure 4**). There was a significant effect of age on the number of oxytocin cells in the PVH (F_2,21_=3.654, p=0.04). Post-hoc test revealed that the number of oxytocin cells increased from 12- to 18-months (q_21_=3.75, p=0.04), while there was a weak trend towards increasing number of cells from 6- to 18-months (q_21_=3.02, p =0.11). Meanwhile, there was no effect of age on the number of vasopressin cells in the PVH (F_2,17_=0.43, p=0.66). Based on these findings, we asked whether there were any correlations between oxytocin cell number and partner directed behaviors. However, correlations between cell counts and partner preference scores or partner huddle time revealed no significant associations on any test day whether examining age groups separately or aggregated (**Supplemental Table 2**).

**Figure 4.**
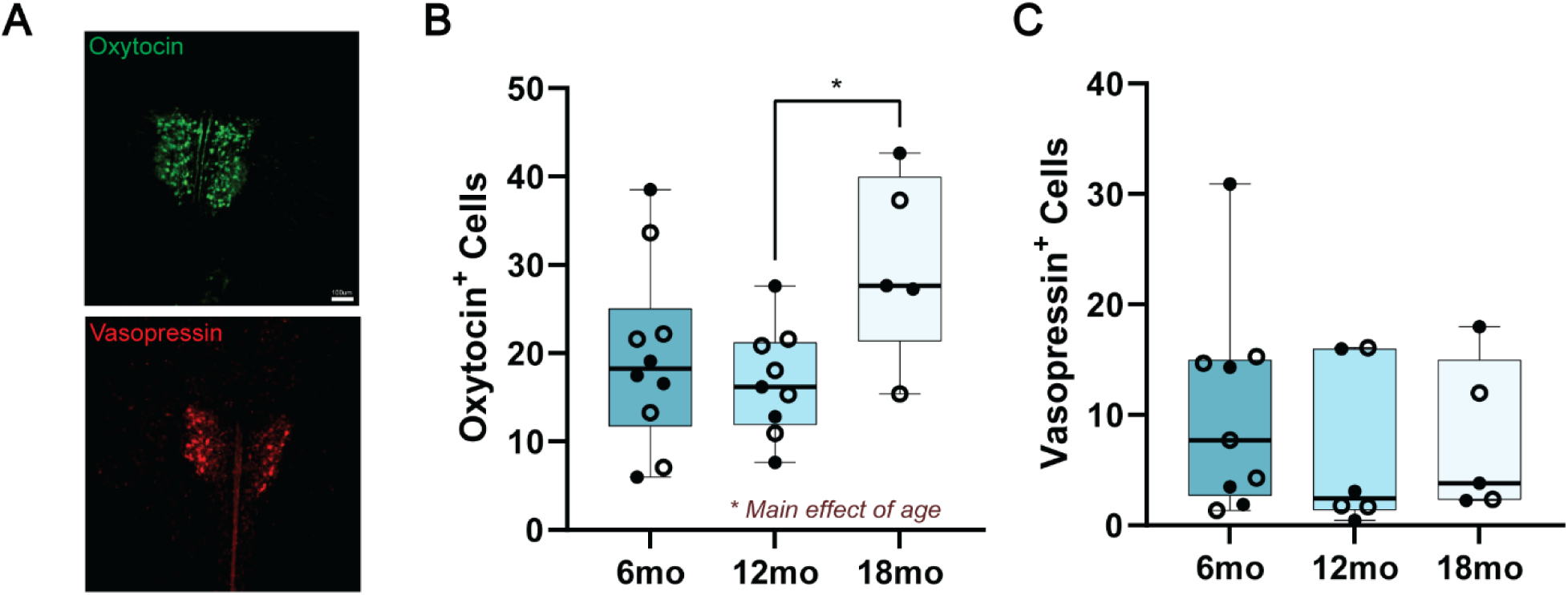
Quantification of oxytocin and vasopressin-producing cells in the PVH reveals a selective effect of aging on the number of oxytocin, but not vasopressin cells. A) Representative images of oxytocin and vasopressin staining in the PVH from a 6-month animal. Scale bar 100µm. B) There was a main effect of age of the number of oxytocin cells in the PVH. Post-hoc tests revealed a significantly greater number of oxytocin cells in 18-month animals compared to 12-month animals. B) Age did not influence the number of vasopressin cells in the PVH. Empty circles = female, closed circles = male. N = 5-10 animals per group; *p<0.05.

## 4. Discussion

Virtually all previous studies interrogating pair bond dynamics in prairie voles have been performed in young adult animals. However, we know from human literature that pair bonds evolve over a life course and are especially important to maintaining health and well-being in late stages of life. Here, we provide the first comprehensive overview of how age sex-dependently impacts pair bonding in a preclinical rodent model, monogamous prairie voles. Specifically, we found that aging accelerates bond formation in males, while aging in females led to an increase in partner directed affiliative behaviors during bond maturation. Furthermore, females appear to be more sensitive to long term separation; unlike males, they displayed lower partner preference scores and reduced partner-directed affiliative behavior after 4 wks separation, which was most prevalent at 12-months. Together, these data provide new evidence on the sex-dependent effects of age on the formation and maintenance of pair bonds, which could be useful for understanding how bonds and partner loss impact emotional well-being during aging.

### 4.1 Patterns of male pair bonds are consistent up to 12-months and then accelerate by 18-months, perhaps reflecting reproductive urgency

We began by confirming previous reports that adult male voles require longer cohabitation to form bonds (Brusman et al., 2021; DeVries and Carter, 1999). While this had previously been demonstrated in animals ∼2-3 months old, our results show that this extends up to at least 12 months of age. Interestingly, at 18-months-old, male voles developed a robust partner preference after only 2 days of cohabitation. By 2 wks, male animals of all ages tested displayed a significant partner preference. We further show that partner preference persists after 4 wks of separation regardless of age in male voles.

Our finding that 18-month-old males form partner preference more quickly is consistent with a prior observation that aged male prairie voles tend to form stronger bonds more quickly than young males regardless of pairing history (Kenkel et al., 2019). Male voles ∼2-3 years old showed a partner preference after short cohabitation times, while younger animals up to ∼1-1.5 years old spent significantly more time in social contact with stranger animals (Kenkel et al., 2019). We posit that these findings may reflect increased reproductive urgency in male animals in response to the real or perceived negative impacts of aging on reproductive performance and success (Comizzoli and Ottinger, 2021). Indeed, aged male animals in other species are less selective concerning mate choices despite potential reproductive disadvantages (Baxter et al., 2020; Churchill et al., 2019; Rundus et al., 2015).

Partner preference phenotypes across aging may also be optimized for the varied mating strategies voles employ in the wild (Getz and McGuire, 1993; Shuster et al., 2019). Male prairie voles can either form a territory (residents) or not (wanderers). Wandering in males typically results in longer survival and greater number of sired offspring, although this may partly depend on factors such as a population density (Getz and McGuire, 1993; Okhovat et al., 2015; Shuster et al., 2019). However, within male voles that adopt a resident strategy, males sire a higher number of litters when in a monogamous partnership (Shuster et al., 2019), for which quicker bond formation and robust maintenance of partner preference may be advantageous in older males.

### 4.2 Female pair bond phenotypes dominate intra-pair behaviors during free interaction

We also confirmed previous reports that young female voles develop a partner preference after only 2 days of cohabitation, which was maintained also at the 2 wk cohabitation timepoint (Brusman et al., 2021; DeVries and Carter, 1999). This was true of all three age groups tested. Unlike our previous report, we only observed increases in partner-directed affiliative behaviors (huddle time) in 18-month female voles at the 2 wk timepoint. The amount of partner-directed huddling was also greater in 18-month animals at 2 wks compared to either other age group. It is important to note that animals from our previous study were only ∼2-3 months old (Brusman et al., 2021), indicating that age has nuanced impacts on the expression of bond phenotypes.

Metrics of partner preference in response to separation had been well described in male voles but was relatively uncatalogued in females. Females display a host of isolation-induced phenotypes, including anhedonia, increased anxiety-like phenotypes and aggression, and neuroendocrine disruption (Grippo et al., 2008, 2007; McNeal et al., 2014; Scotti et al., 2015; Watanasriyakul et al., 2022). One study reported persistence of pair bonds following 4 wks separation, without any effect of sex in ∼2-6 month old voles (Fricker et al., 2023). In our experiments, partner preference after 4 wks separation followed a U-curve, where 6- and 18-month females demonstrated a strong partner preference, but 12-month animals did not. However, across all ages at the 4 wks separation timepoint, there was a tendency for females to decrease huddling time with a partner. These phenotypes may reflect an overall change in social behavior, as aging in unpaired voles reduces both prosocial and aggressive behaviors (Powell et al., 2022). However, within the context of a pair bond, these shifts in behavior could promote flexibility to engage with novel animals and potentially form new bonds. Interestingly, about half of the 12-month female voles displayed a partner preference after 4 wks separation while the other half did not, a split that was not observed in any other group. If such trends are seen in future groups of animals, they could provide a way to investigate the neural mechanisms supporting this switch in preference across ages. Furthermore, we cautiously suggest that this flexibility in females is not due to the negative effects of aging on reproductive success since there were similar rates of pregnancy in all age groups (data not shown). We did not, however, measure other signs of reproductive fitness, such as litter size, that may also explain such a U-shape curve in response to partner loss in females (Curtis, 2010; Resendez et al., 2016).

Similar to males, females can adopt wandering or resident phenotypes to boost reproductive success, which may also be partially reflected in our PPT outcomes (Shuster et al., 2019). Specifically, wandering females tend to produce more offspring when in polyandrous versus monandrous partnerships (Shuster et al., 2019), which is what might be expected in 12-month females following the loss of a pair bonded partner. The age-dependency for the adoption of either strategy in females is an open question, but does raise the interesting possibility that voles coordinate their mating tactics at a given age in order to maximize reproductive fitness.

Consistent with previous findings from our lab, we show that bond phenotypes of female animals appear to dominate joint behavior expressed during free interaction tests (Brusman et al., 2021). This is most apparent in 12-month pairs after 4 wks separation, where 12-month females lacked a partner preference, which was paralleled by decreased in close social contact during free interaction, suggesting that females rather than males determine overall interaction levels. This lack of partner preference in females does not appear to be associated with male preference on the same test day, as nearly all males showed a robust partner preference.

### 4.3 Initial interactions during PPT reflect sex-specific differences in partner preference

Investigation of the initial 5 minutes and subsequent 15 minutes of partner preference tests allowed us to assess novelty and habituation of interactions with partners and stranger conspecifics at each age. In general, males sampled both partner and stranger animals during both time periods analyzed. This agrees with a previous report showing that bonded males will equally press levers for access to partner or stranger female animals (Brusman et al., 2021; Vahaba et al., 2022). The exception was following 4 wks separation in 12- and 18-month male animals, which displayed a strong partner preference during the 5-20 minute time period. This result suggests that age and test day could modulate behavior regarding social motivation. As such, it would be interesting to extend these operant tasks to different ages and test days.

Meanwhile, females displayed a greater likelihood for partner preference during both time periods, except for 6-month animals which preferred stranger animals in the very early stages of bonding (2 days cohabitation) in the first 5 minutes. Unlike males, these preferences are largely indicative of preference over the entire 3-hour PPT. This is also in agreement with operant literature showing female animals will preferentially press levers for partner over stranger male animals (Vahaba et al., 2022).

### 4.4 Neurochemical systems influencing pair bond dynamics across aging

We found that the number of oxytocin, but not vasopressin, cells in the PVH increased with age. This increase was specific when comparing 12- and 18-month animals, while a weak trend was noted between 6- and 18-month animals. While we were underpowered to assess any sex differences in number of oxytocin or vasopressin cells over the course of aging, there were no obvious trends which matches previous vole and human literature (Fricker et al., 2023; Ishunina and Swaab, 1999; Wierda et al., 1991). Data from human literature show no changes in oxytocin cell number and some changes in composition of vasopressin-positive cells over the course of aging or in AD (Fliers et al., 1985; Ishunina and Swaab, 1999, 1999; Wierda et al., 1991). We offer two possibilities for these discrepancies. The first is that there are species differences between voles and humans. The other, non-mutually exclusive explanation is that natural variability in form and function of these socially responsive neural systems lead to different behavioral outputs, even within a specific age range (Audunsdottir and Quintana, 2022; Blumenthal and Young, 2023; Ebner et al., 2013). While there were no correlations between oxytocin cell numbers and partner preference measurements at any age or test day, it’s possible that other metrics (receptor density, innervation patterns, release, and activity) would better predict behavioral performance. For example, natural variation in oxytocin receptor density in the nucleus accumbens can predict partner preference and mating strategies (King et al., 2016; Ophir et al., 2012), and variation in vasopressin receptor levels in the retrosplenial cortex reflect mating strategies in the wild (Okhovat et al., 2015). Yet how this and other variations change with age is unknown. It is also possible that some of the changes observed here are not attributable to age alone but rather reflect age-dependent response to loss of a bonded partner, given that tissue was taken after 4 wks separation (Fricker et al., 2023; Grippo et al., 2007). For example, greater numbers of oxytocin cells in the PVH may support the maintenance of partner preference after separation in 18-month males, while lower cell numbers may promote flexibility seen in 12-month females.

## 5. Conclusions

Our study is the first to show that age has sex-specific effects on the presentation of pair bond behaviors in prairie voles. Specifically, that aging accelerates pair bond formation in males while response to partner loss in females appears to follow a U-shaped curve that dominates expression of intra-pair dynamics during unconstrained free interaction. These behaviors are likely influenced by changes in form and function of socially responsive brain regions, such as the observed increase in oxytocin cell number of the PVH during aging. These findings raise the possibility of general alterations to other social behaviors that can be systematically evaluated in prairie voles such as aggression, emotional contagion, and consoling behaviors. This work could be extended to investigate how aging affects non-social phenotypes in response to partner loss. Overall, this work establishes the monogamous prairie vole as a useful preclinical model for studying the effects of normal and abnormal aging on attachment-related behaviors.

## Funding and Acknowledgements

This work was supported by funding from the National Institute of Mental Health (R01 MH125423), National Institute of Aging (U24 AG072701), and National Science Foundation (IOS-2045348) all awarded to ZRD.

## CRediT authorship contribution statement

**Michael A. Kelberman:** Conceptualization, Formal Analysis, Investigation, Data Curation, Writing – Original Draft, Writing – Review and Editing, Visualization **Kelly Winther:** Investigation, Data Curation, Writing – Review and Editing, Visualization **Yana M:** Investigation **Zoe Donaldson:** Conceptualization, Writing – Review and Editing, Supervision, Funding Acquisition

---

**Supplemental Figure 1.**
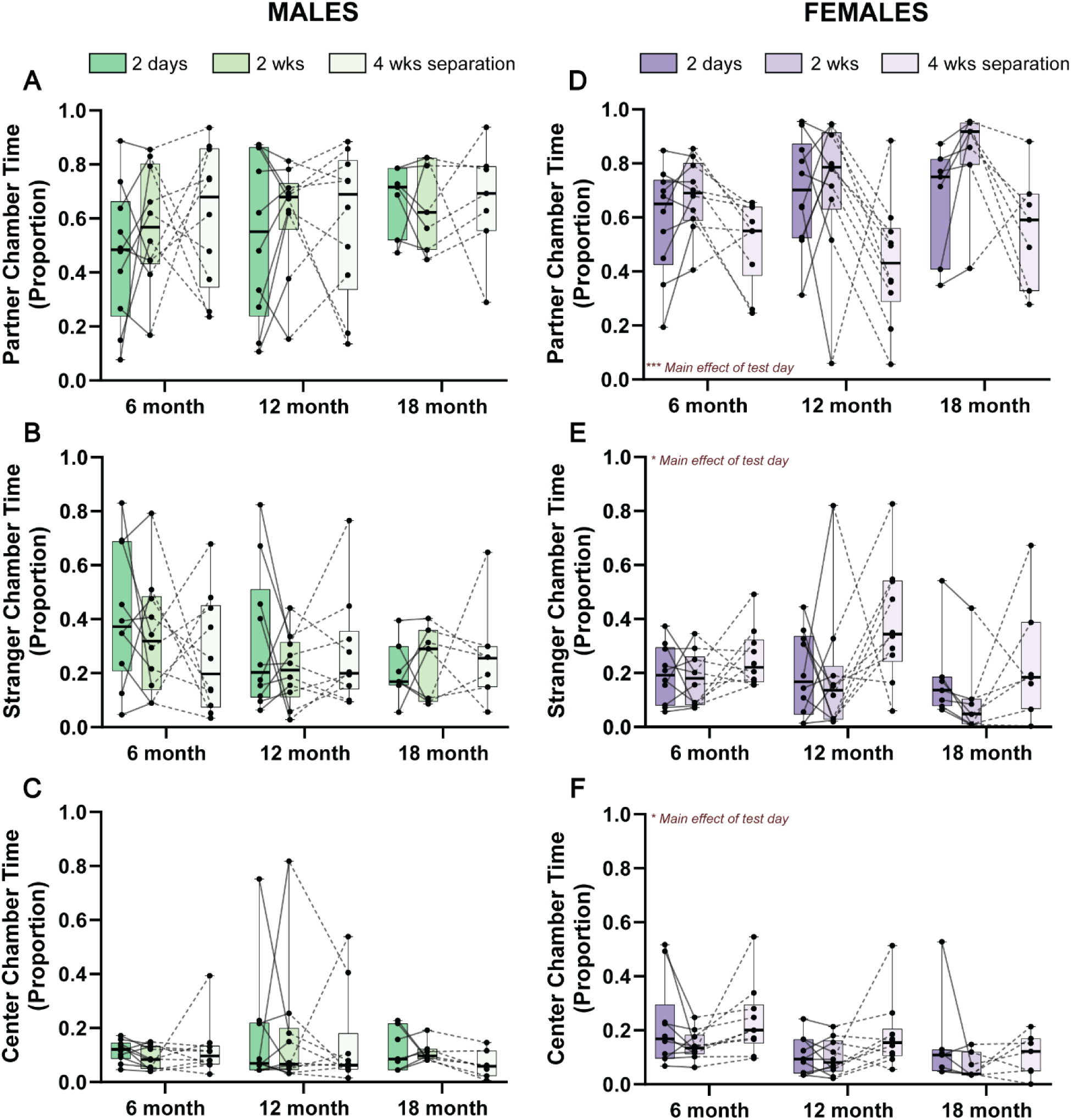
Effects of age and test day on proportion of time animals spent in different areas of the partner preference arena. A) Males across age and test day do not spend a greater or less proportion of their time in the partner B) stranger C) or center chamber of the partner preference arena. D) Female animals spent a greater proportion of time in the partner chamber after 2 wks cohabitation and less after 4 wks separation. E) These effects were mirrored in the opposite direction when observing proportion of time spent in the stranger chamber. F) With increasing age, there was a gradually decreasing proportion of time spent in the center chamber during the partner preference task in female animals. N=7-10 animals per group; *p<0.05, ***p<0.001.

**Supplemental Table 1:** 12-month Free Interaction 15 Minute Time Bin Statistics.

| 12-months |  |  |  |  |
| --- | --- | --- | --- | --- |
| Time Bin | Test | q statistic | df | p value |
| 1 | 2 days vs. 2 wks | 1.777 | 198 | 0.4215 |
|  | 2 days vs. 4 wks sep | 2.851 | 198 | 0.1110 |
|  | 2 wks vs. 4 wks sep | 1.074 | 198 | 0.7284 |
| 2 | 2 days vs. 2 wks | 0.9542 | 198 | 0.7785 |
|  | 2 days vs. 4 wks sep | 2.370 | 198 | 0.2170 |
|  | 2 wks vs. 4 wks sep | 1.416 | 198 | 0.5769 |
| 3 | 2 days vs. 2 wks | 1.128 | 198 | 0.7049 |
|  | 2 days vs. 4 wks sep | 3.121 | 198 | 0.0725 |
|  | **2 wks vs. 4 wks sep** | **4.249** | **198** | **0.0084** |
| 4 | 2 days vs. 2 wks | 1.188 | 198 | 0.6787 |
|  | 2 days vs. 4 wks sep | 3.162 | 198 | 0.0677 |
|  | 2 wks vs. 4 wks sep | 1.975 | 198 | 0.3448 |
| 5 | 2 days vs. 2 wks | 1.808 | 198 | 0.4091 |
|  | **2 days vs. 4 wks sep** | **5.212** | **198** | **0.0009** |
|  | **2 wks vs. 4 wks sep** | **3.404** | **198** | **0.0446** |
| 6 | 2 days vs. 2 wks | 1.491 | 198 | 0.5436 |
|  | **2 days vs. 4 wks sep** | **3.970** | **198** | **0.0151** |
|  | 2 wks vs. 4 wks sep | 2.478 | 198 | 0.1885 |
| 7 | 2 days vs. 2 wks | 2.392 | 198 | 0.2110 |
|  | **2 days vs. 4 wks sep** | **5.575** | **198** | **0.0003** |
|  | 2 wks vs. 4 wks sep | 3.183 | 198 | 0.0654 |
| 8 | 2 days vs. 2 wks | 1.784 | 198 | 0.4186 |
|  | **2 days vs. 4 wks sep** | **5.514** | **198** | **0.0004** |
|  | **2 wks vs. 4 wks sep** | **3.730** | **198** | **0.0244** |
| 9 | **2 days vs. 2 wks** | **5.000** | **198** | **0.0015** |
|  | **2 days vs. 4 wks sep** | **9.138** | **198** | **<0.0001** |
|  | **2 wks vs. 4 wks sep** | **4.139** | **198** | **0.0106** |
| 10 | **2 days vs. 2 wks** | **5.244** | **198** | **0.0008** |
|  | **2 days vs. 4 wks sep** | **10.22** | **198** | **<0.0001** |
|  | **2 wks vs. 4 wks sep** | **4.974** | **198** | **0.0016** |
| 11 | 2 days vs. 2 wks | 1.426 | 198 | 0.5725 |
|  | **2 days vs. 4 wks sep** | **8.550** | **198** | **<0.0001** |
|  | **2 wks vs. 4 wks sep** | **7.124** | **198** | **<0.0001** |
| 12 | 2 days vs. 2 wks | 0.2429 | 198 | 0.9839 |
|  | **2 days vs. 4 wks sep** | **9.268** | **198** | **<0.0001** |
|  | **2 wks vs. 4 wks sep** | **9.511** | **198** | **<0.0001** |

**Supplemental Table 2:**
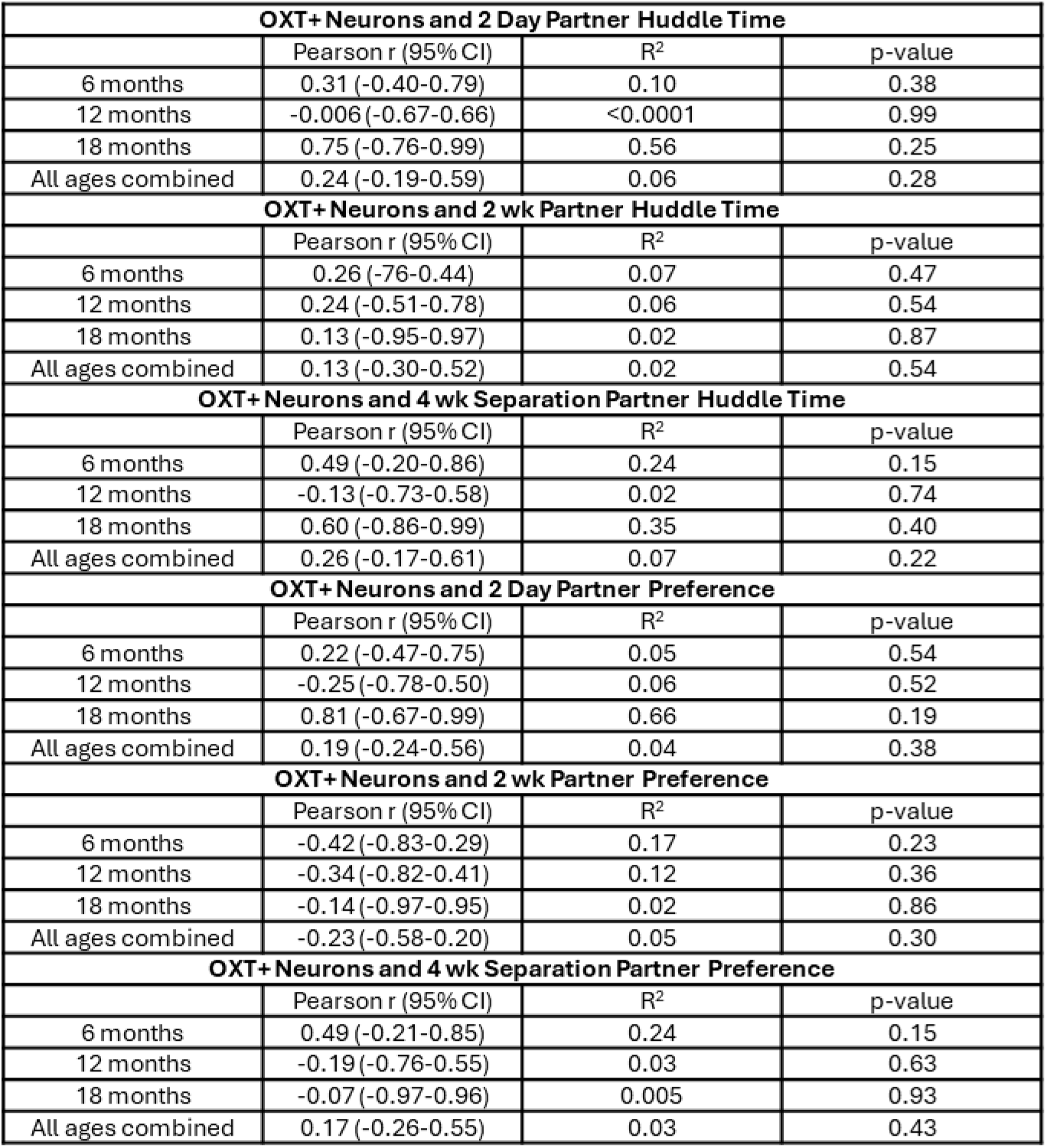
Correlation Statistics Between PPT Metrics and Oxytocin Cell Counts.

| **OXT+ Neurons and 2 Day Partner Huddle Time** |  |  |  |
| --- | --- | --- | --- |
|  | Pearson r (95% CI) | R<sup>2</sup> | p-value |
| 6 months | 0.31 (-0.40-0.79) | 0.10 | 0.38 |
| 12 months | -0.006 (-0.67-0.66) | <0.0001 | 0.99 |
| 18 months | 0.75 (-0.76-0.99) | 0.56 | 0.25 |
| All ages combined | 0.24 (-0.19-0.59) | 0.06 | 0.28 |
| **OXT+ Neurons and 2 wk Partner Huddle Time** |  |  |  |
|  | Pearson r (95% CI) | R<sup>2</sup> | p-value |
| 6 months | 0.26 (-0.76-0.44) | 0.07 | 0.47 |
| 12 months | 0.24 (-0.51-0.78) | 0.06 | 0.54 |
| 18 months | 0.13 (-0.95-0.97) | 0.02 | 0.87 |
| All ages combined | 0.13 (-0.30-0.52) | 0.02 | 0.54 |
| **OXT+ Neurons and 4 wk Separation Partner Huddle Time** |  |  |  |
|  | Pearson r (95% CI) | R<sup>2</sup> | p-value |
| 6 months | 0.49 (-0.20-0.86) | 0.24 | 0.15 |
| 12 months | -0.13 (-0.73-0.58) | 0.02 | 0.74 |
| 18 months | 0.60 (-0.86-0.99) | 0.35 | 0.40 |
| All ages combined | 0.26 (-0.17-0.61) | 0.07 | 0.22 |
| **OXT+ Neurons and 2 Day Partner Preference** |  |  |  |
|  | Pearson r (95% CI) | R<sup>2</sup> | p-value |
| 6 months | 0.22 (-0.47-0.75) | 0.05 | 0.54 |
| 12 months | -0.25 (-0.78-0.50) | 0.06 | 0.52 |
| 18 months | 0.81 (-0.67-0.99) | 0.66 | 0.19 |
| All ages combined | 0.19 (-0.24-0.56) | 0.04 | 0.38 |
| **OXT+ Neurons and 2 wk Partner Preference** |  |  |  |
|  | Pearson r (95% CI) | R<sup>2</sup> | p-value |
| 6 months | -0.42 (-0.83-0.29) | 0.17 | 0.23 |
| 12 months | -0.34 (-0.82-0.41) | 0.12 | 0.36 |
| 18 months | -0.14 (-0.97-0.95) | 0.02 | 0.86 |
| All ages combined | -0.23 (-0.58-0.20) | 0.05 | 0.30 |
| **OXT+ Neurons and 4 wk Separation Partner Preference** |  |  |  |
|  | Pearson r (95% CI) | R<sup>2</sup> | p-value |
| 6 months | 0.49 (-0.21-0.85) | 0.24 | 0.15 |
| 12 months | -0.19 (-0.76-0.55) | 0.03 | 0.63 |
| 18 months | -0.07 (-0.97-0.96) | 0.005 | 0.93 |
| All ages combined | 0.17 (-0.26-0.55) | 0.03 | 0.43 |

