## Supplementary figures and images for "Aging leads to sex-dependent effects on pair bonding and increased number of oxytocin-producing neurons in monogamous prairie voles"

### Supplemental Figure 1

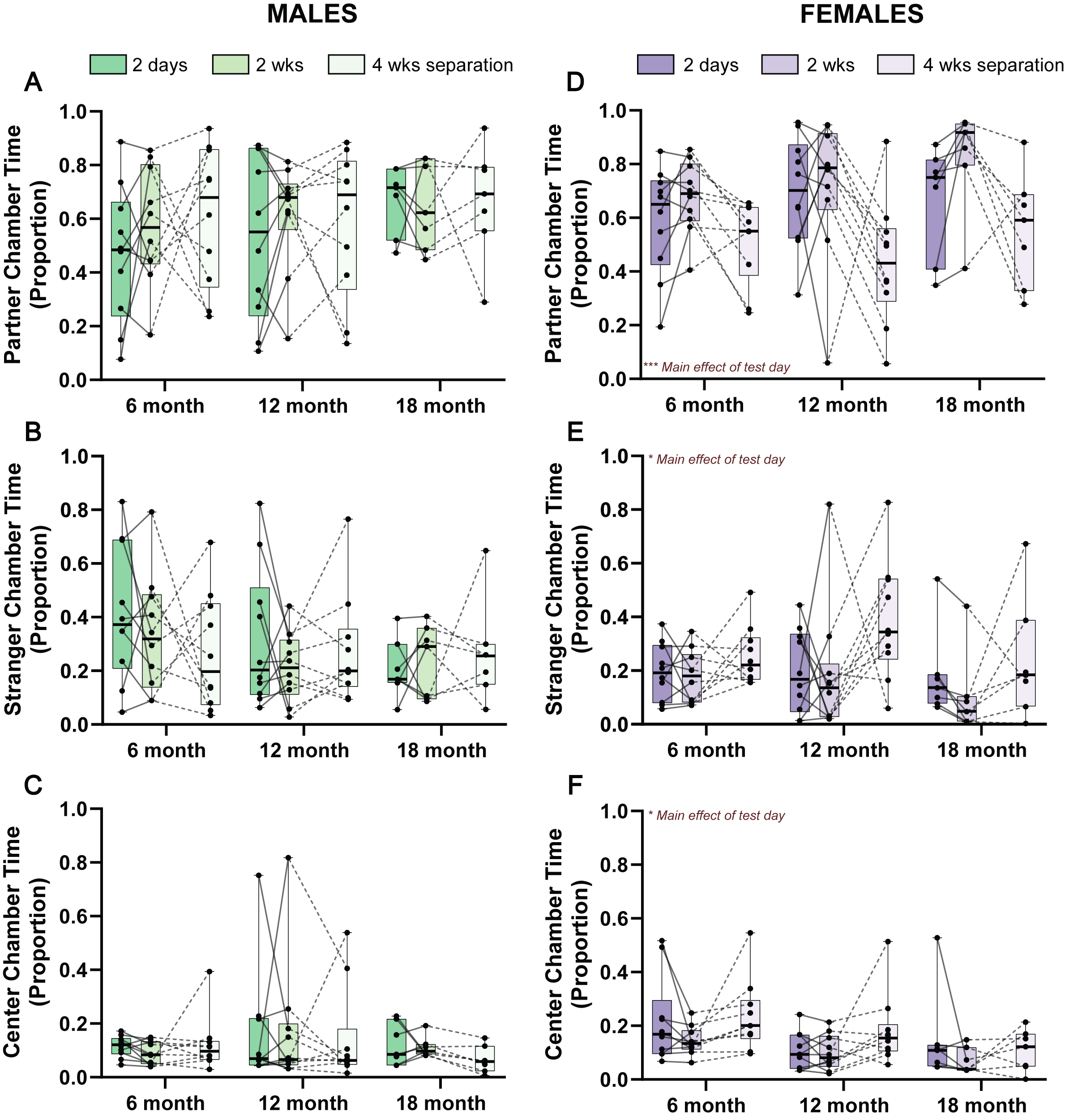
